# Oxygen-adaptive plasticity of Asgard archaea dependent on terminal oxidase and globin

**DOI:** 10.1101/2025.11.07.685452

**Authors:** Zhongyi Lu, Runyue Xia, Aoxiang Xu, Jiazheng Gu, Huifang Cai, Yang Liu, Eugene V. Koonin, Meng Li

## Abstract

The oxygenation of ancient Earth is thought to have driven eukaryogenesis, beginning with the endosymbiosis of an aerobic alphaproteobacterium (proto-mitochondria) with an archaeal host. Given that the archaeal host likely evolved from within Asgard archaea (phylum *Promethearchaeota*), the metabolic traits of Asgard archaea could provide key insights into eukaryotic origins. Although Asgard archaea cultured to date are obligate anaerobes, their genomes encode oxygen-adaptive proteins, suggesting they might be oxygen-tolerant. Here, we demonstrate that some Asgard archaea, in particular, *Hodarchaeales*, the closest known relatives of eukaryotes, and *Kariarchaeaceae*, exhibit oxygen adaptation mediated by terminal oxidase and globin. Phylogenetic analysis reveals long-term vertical evolution of terminal oxidases in Asgard archaea, suggesting ancient adaptation to molecular oxygen. By contrast, globin was likely acquired by Asgard archaea via horizontal gene transfer from facultative aerobic *Chloroflexales* bacteria. Heterologous expression of the Asgard globin enhances aerobic growth of *Haloarchaea* and *Escherichia coli* in the presence of terminal oxidase-dependent electron transfer chain, suggesting that Asgard growth benefits from ambient oxygen. The Asgard globin gene is embedded in an oxygen-sensitive bidirectional promoter region, with one promoter driving oxygen-induced globin expression, and the other anaerobically activating expression of two enzymes, PdxS and PdxT, involved in a pyridoxal 5’-phosphate biosynthesis. The Asgard globin and promoter region exhibit high functional robustness across archaea and bacteria, and could contribute to the symbiosis between the Asgard and aerobic bacterial partners. These findings highlight the oxygen-adaptive plasticity of Asgard archaea and its potential contribution to eukaryogenesis.

## INTRODUCTION

The Great Oxidation Event fundamentally shaped the evolution of life on Earth ^1–4^. The subsequent rise of aerobic metabolism enabled high-energy flux of prokaryotic cells, and drove their genome expansion and increased cellular complexity, the eukaryotic hallmarks ^5,6^. However, excess molecular oxygen is toxic to cells, and therefore, organisms evolve adaptive mechanisms that detoxify and utilize oxygen ^7,8^. Among these mechanisms, terminal oxidases, also known as heme-copper oxidases, catalytically reduce oxygen to water ^9^. The terminal oxidases can be functionally divided into cytochrome *c* oxidases (Cox) and ubiquinol oxidases (Uox). Although Cox and Uox are homologous, they substantially differ in their distribution across biological diversity and their molecular mechanisms ^10^. Cox is widely distributed in aerobic bacteria and is a key component of the mitochondrial electron transport chain in eukaryotes where it functions as the terminal oxidase, accepting electrons from cytochrome *c* ^11^. By contrast, Uox accepts electron from ubiquinol and is required for oxidative stress resistance across bacteria and archaea ^11^. Noteworthy, certain Cox accepts electrons from both cytochrome c and ubiquinol, indicative of functional versatility of the terminal oxidases in utilizing widely different substrates ^12^.

Globins play crucial roles in oxygen adaption via oxygen binding, transport, and storage across the tree of life ^13–16^. The globin superfamily includes bacterial truncated hemoglobin (trHbO), vertebrate hemoglobin, and archaeal protoglobin all of which share structurally conserved heme-containing 2-over-2 or 3-over-3 alpha-helical sandwich fold ^17–19^. In *Mycobacterium* and *Vitreoscilla*, trHbO expression is upregulated under hypoxic condition, promoting aerobic respiration by triggering terminal oxidase ^16,20,21^.

Beyond the terminal oxidase and hemoglobin, the prokaryotic pyridoxal 5’-phosphate (PLP, active form of vitamin B6) biosynthesis pathway is functionally associated with oxygen because some of the PLP-dependent enzymes utilize oxygen as a co-substrate ^22,23^. There are two variants of the PLP pathway, deoxyxylulose 5-phosphate (Dxp)-dependent and Dxp-independent ones. The Dxp-dependent pathway has a relatively narrow phylogenetic distribution, being scattered among several bacterial clades, whereas the Dxp-independent pathway is widespread across archaea, bacteria, plants, and fungi ^24,25^. The Dxp-independent pathway relies on two enzymes, pyridoxal 5’-phosphate synthase (PdxS) and pyridoxal 5’-phosphate synthase glutaminase (PdxT), which jointly catalyze the conversion of glutamine to PLP ^24^. Despite its broad occurrence, the molecular mechanism and regulation of the Dxp-independent pathway remain poorly characterized.

The oxygen-adaptive systems enable some microorganisms to survive across a broad range of oxygen concentrations. For example, nanoxia is characterized by extremely low, nanomolar oxygen concentrations, occurring in habitats such as marine oxygen minimum zone, aquatic sediments, wetland soils, and animal guts ^26,27^. Critically, unlike true anoxia, the nanoxic environments experience constant, even if low, oxygen influx from adjacent oxic zones, resulting in highly dynamic fluctuations in local oxygen levels. The microorganisms inhabiting these niches, for example, the bacterium *Bacteroides fragilis* and the archaeon *Thermococcus onnurineus*, are typically considered obligate anaerobes because their growth is inhibited by oxygen concentration >1 μM ^28,29^, Nevertheless, their growth can be stimulated by nanomolar concentration of oxygen, seemingly owing to their terminal oxidases that catalyze oxygen reduction to water ^27,28^. Despite these findings, the molecular and metabolic mechanisms behind the paradoxical stimulation of the growth of nanoxic microbes by oxygen remain poorly characterized.

Under the current leading scenario of eukaryogenesis, eukaryotes evolved as a result of endosymbiosis, whereby an archaeal host cell engulfed an alphaproteobacterial ancestor of the mitochondria ^30–32^. Given that the mitochondrial ancestor possessed the aerobic electron transport chain, the archaeal host most likely evolved oxygen adaptation mechanisms ^4,33^. The archaeal ancestor of eukaryotes is believed to originate from within Asgard archaea (phylum *Promethearchaeota*), most likely, as the sister group of the order *Hodarchaeales* within the class *Heimdallarchaeia* ^34–39^. Metabolic reconstruction suggests that several Asgard lineages within *Heimdallarchaeia*, including *Hodarchaeales*, *Gerdarchaeales*, *Kariarchaeaceae*, and *Heimdallarchaeaceae*, are facultative aerobic heterotrophs ^37,40,41^. Consistent with this inference, recent metagenomic analyses showed that *Heimdallarchaeia* encode homologues of electron transport chain complexes including terminal oxidase and globin, which could be involved in aerobic respiration ^42^. However, in contrast to these metagenomic analyses, most recent isolation of members of *Hodarchaeales* shows a strictly anaerobic lifestyle, suggesting that the aerobic respiration chain is not active in these organisms^43^. These findings make Asgard adaptation to environmental oxygen a particularly intriguing problem.

Here, we identify and characterize terminal oxidase and globin in three groups of Asgard archaea, *Kariarchaeaceae*, *Gerdarchaeales*, and *Hodarchaeales*, and their homologs in TACK archaea and Haloarchaea. Phylogenetic, genetic, and biochemical analyses of these proteins reveal oxygen-adaptive flexibility of Asgard archaea dependent on the regulation of terminal oxidases and globin.

## RESULTS

### Origin and evolution of terminal oxidases in Asgard archaea

It has been reported previously that Asgard archaea encode some terminal oxidase subunits ^42,43^. To investigate the spread and evolutionary history of these proteins, we screened our local database of Asgard genomes ^37^, and identified a set of terminal oxidase subunits in *Kariarchaeaceae*, *Gerdarchaeales*, and *Hodarchaeales* within *Heimdallarchaeia*. In Asgard genomes, genes encoding the terminal oxidase subunits comprise a predicted operon which in *Gerdarchaeales* and *Hodarchaeales* includes the genes encoding CyoA, CyoB, CyoC, and CyoD (Fig. 1a). These subunits constitute a typical Uox complex, exemplified by that of *Escherichia coli* (Figure 1b), where CyoA contains a peripheral domain with binuclear copper-binding site, whereas CyoB harbors a heme-copper-dependent catalytic center ^10,44^. In contrast to these typical Uox, *Kariarchaeaceae* exhibits a unique operon organization, retaining CyoC and CyoD (although absent in some genomes) and featuring a distinct CyoB variant, with an N-terminal extension homologous to CyoC. To further characterize these Asgard terminal oxidases, we modeled terminal oxidase structures of *Hodarchaeales* and *Kariarchaeaceae* using AlphaFold3 ^45^, and found that the predicted structure of the terminal oxidase of *Hodarchaeales* and *Kariarchaeaceae* closely resembled that of the *E. coli* Uox (root-mean-square deviation, RSMD: 0.890 Å and 1.421 Å, respectively), with the peripheral region of Asgard CyoA interacting with CyoB (Figure 1b). Despite the distinct CyoB architecture, the *Kariarchaeaceae* terminal oxidase was also found to be closely structurally similar to the counterpart from *Hodarchaeales* (RSMD: 0.732 Å), demonstrating conservation of terminal oxidases in *Heimdalarchaeia*. Collectively, these findings demonstrate the broad presence of terminal oxidases in Asgard archaea, especially, in *Heimdalarchaeia*, with lineage-specific variations in subunit composition and architecture.

**Figure 1.**
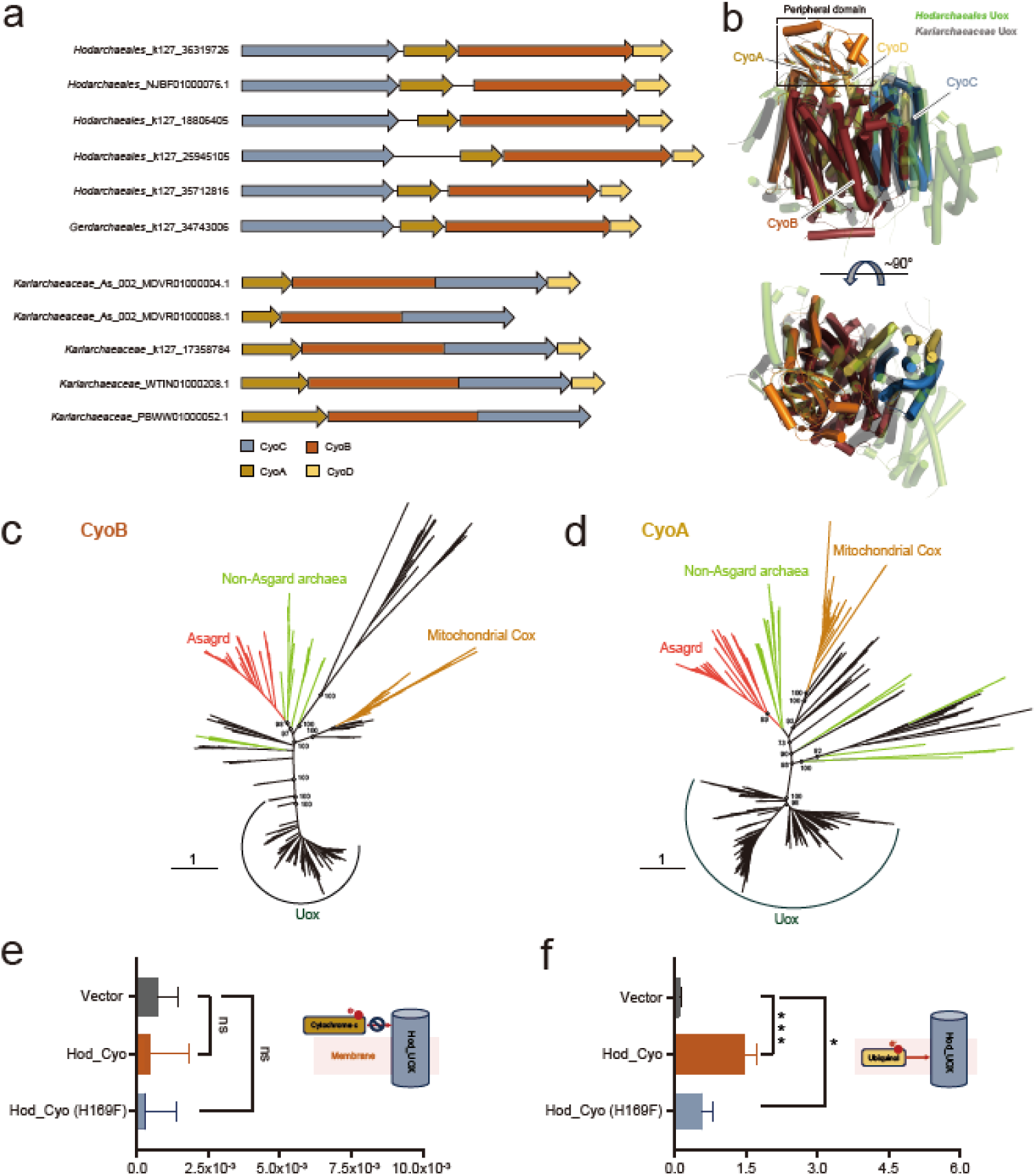
Terminal oxidases in Asgard archaea. (a) Operon organization of the genes encoding terminal oxidases in *Hodarchaeales*, *Gerdarchaeales*, and *Kariarchaeaceae*. The N-terminal extension homologous to CyoC of *Kariarchaeaceae* CyoB variant was labeled. (b) Structural alignment of the terminal oxidases of *Hodarchaeales* (green), *Kariarchaeaceae* (grey), and *E. coli* (PDB number: 6WTI). The *E. coli* terminal oxidases subunits are highlighted in distinct colors (CyoA, orange; CyoB, red; CyoC, blue; CyoD, yellow) and the peripheral domain is labeled. (c) Unrooted maximum likelihood tree of CyoB homologs. The tree was inferred from an alignment of 287 sequences across Bacteria (191 sequences from *Bacillati* and *Pseudomonadota*, black), eukaryotic mitochondria (23 sequences, yellow), Asgardarchaeota (50 sequences, red), and non-Asgard archaea (23 sequences from TACK and Euryarchaeota; green) based on best-fit model LG+F+I+G4 in IQ-Tree. Bootstrap values are shown for key nodes. (d) Unrooted maximum likelihood tree of CyoA homologs. The tree was inferred from an alignment of 328 sequences across Bacteria (218 sequences from *Bacillati* and *Pseudomonadota*, black), eukaryotic mitochondria (32 sequences, yellow), *Asgardarchaeota* (49 sequences, red), and non-Asgard archaea (29 sequences from TACK and Euryarchaeota; green) based on best-fit model Q.pfam+G4 in IQ-Tree. Bootstrap values are shown for some nodes. (e-f) The oxidase activity of Asgard UOX when cytochrome c (e) and ubiquinone (f) are used as the electron donor. Asterisks indicate significant differences: *p<0.05, ***p<0.001, ns denotes non-significant differences.

To explore the evolution of the terminal oxidases in Asgard archaea, and considering the pivotal role of CyoB as a core subunit, we performed a phylogenetic analysis of CyoB from *Kariarchaeaceae* and *Hodarchaeales* (including recently predicted Asgard Cox1 subunits ^42^), along with CyoB-like proteins from TACK archaea, Euryarchaeota, and mitochondria as well as bacterial CyoB homologs. In the unrooted maximum likelihood phylogenetic tree, the Asgard proteins formed a strongly supported clade, with TACK and Euryarchaeota proteins occupying basal positions, while still grouping within the Cox branch, separately from bacterial and mitochondrial homologs (Figure 1d). We also constructed a phylogenetic tree of CyoA homologs. Consistent with the CyoB phylogeny, Asgard CyoA proteins similarly formed a well-supported clade together with TACK homologs in the Cox branch, while separated from the bacterial and mitochondrial Cox subunits (Figure 1d). Other archaeal CyoA proteins were interspersed with bacterial homologs, likely reflecting multiple horizontal gene transfers. Examination of the sequence alignment shows that the Asgard CyoB subunits encompass bacterial type A CyoB-like motifs with D- and K- proton channels, and histidine ligands for heme and Cu_B_ (Figure S1) ^9,11^. The Cu_A_-binding histidine ligands are conserved between the Asgard and bacterial type A CyoA subunits ^9^. Together, these findings suggest long-term vertical evolution of active terminal oxidases in Asgard archaea.

To characterize an Asgard archaeal terminal oxidase (*Hodarchaeales*_NJBF01000076.1 *cyo*ABCD) biochemically, codon-optimized *cyoA* and *cyoD* were cloned into pCDF-Duet, and *cyoB* and *cyoC* were cloned into pRSF-Duet. Both plasmids were co-transformed into *E. coli* BL21(DE3) for expression. Membrane fractions containing the recombinant oxidase were isolated and assayed for activity using cytochrome c or ubiquinone as electron donors. A loss-of-function mutant (CyoB-H169F), with the conserved Cu_b_-binding histidine replaced by phenylalanine, served as a negative control (Figure S1). As shown in Figure 1e, neither the wild-type *Hodarchaeales* oxidase nor the H169F mutant exhibited detectable activity with cytochrome c. By contrast, robust oxidase activity was observed for the wild-type enzyme with ubiquinone. The H169F mutant showed reduced activity with ubiquinone, the residual activity apparently due to the native ubiquinol oxidase of *E. coli* ^46^. Collectively, these results demonstrate that the Asgard terminal oxidase functions as a ubiquinol oxidase.

The phylogenies of CyoB and CyoA support the monophyly of archaeal A-type Uox, apart from some apparent horizontal transfers of CyoA. In TACK archaea and Haloarchaea, the terminal oxidases are thought to reduce oxygen via electrons from quinols, which are required for aerobic and possibly facultative anaerobic respiration ^47^. By analogy, the Asgard terminal oxidases likely represent an adaptation for the reduction of environmental oxygen.

### Bacterial origin of globin in Asgard archaea

Because globin genes have been identified in Asgard archaeal genomes ^43^, and given their functional connection to terminal oxidase, we systematically searched our Asgard genomic database ^37^ and identified four globins (COG011180) in *Kariarchaeaceae* and one in *Hodarchaeales*. HHpred analysis demonstrates highly significant similarity of the entire sequences of the Asgard globins to trHbO (e value <10^-7^). TrHbO is a truncated hemoglobin that binds oxygen via its iron-dependent heme group and can facilitate bacterial aerobic growth through direct interaction with CyoB of terminal oxidase ^16,20,21^. Notably, two of *Kariarchaeaceae* globin genes (As_030-p_00404 and As_140-p_00992) are adjacent to genes encoding a Serine-tRNA ligase, PdxT, and PdxS. The latter two enzymes constitute the complete Dxp-independent PLP biosynthesis pathway (Fig. 2a).

**Figure 2.**
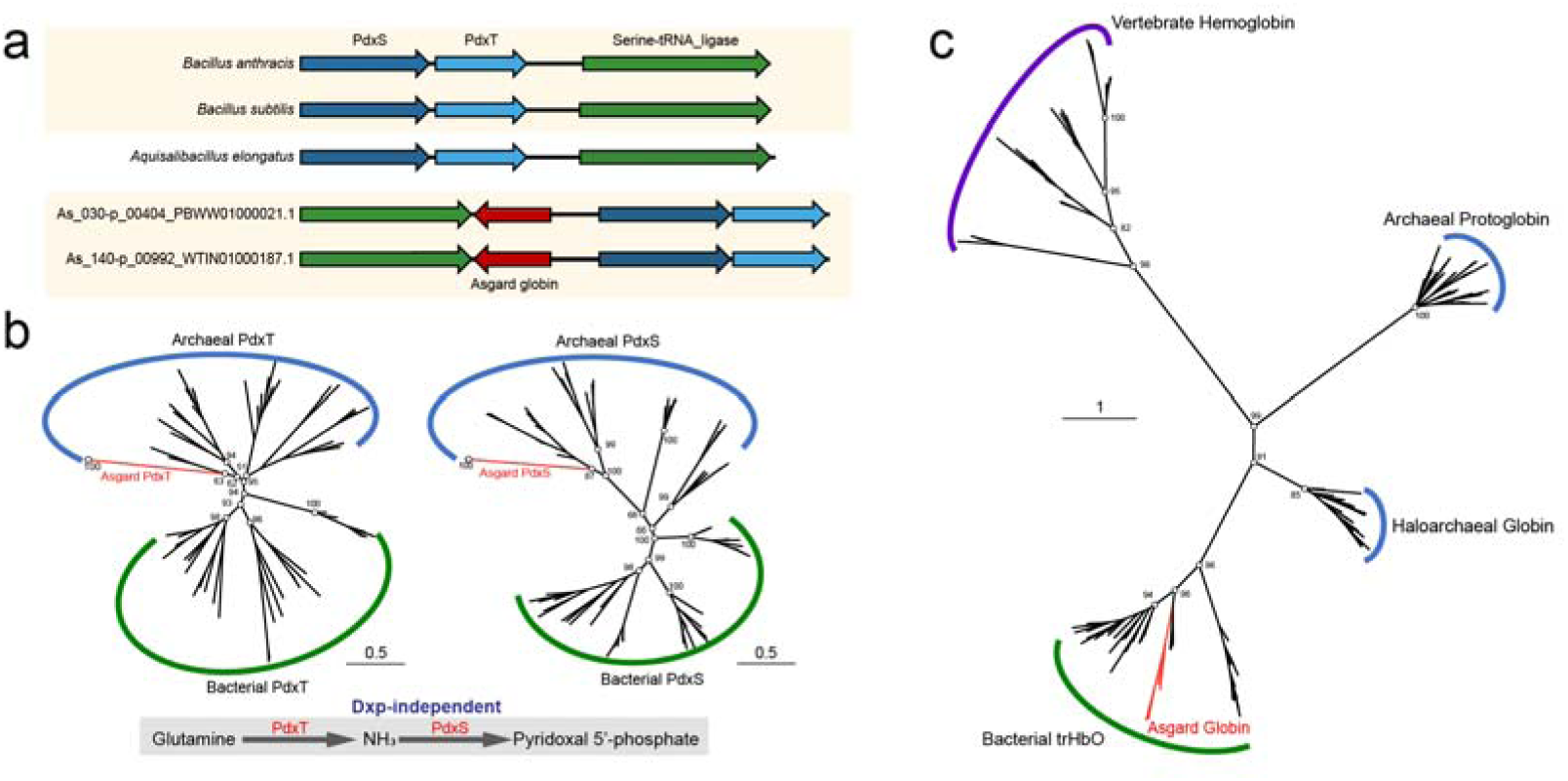
Genomic context and phylogenies of Asgard globins and pyridoxal phosphate biosynthesis enzymes. (a) Globin gene context in *Kariarchaeaceae* and bacteria. Gene names or brief descriptions are provided above or below the corresponding arrows. Homologous genes are color matched across lineages. (b) Unrooted maximum likelihood tree of PdxT (left, best-fit model: LG+I+G4) and PdxS (right, best-fit model: LG+I+G4) from Asgard archaea (red branches), non-Asgard archaea, and bacteria. Bootstrap values are shown for key nodes. The complete deoxyxylulose 5-phosphate (Dxp)-independent pyridoxal 5’-phosphate (PLP) biosynthesis pathway and the involved enzymes PdxT and PdxS are shown. (c) Unrooted maximum likelihood tree of globin superfamily including Asgard globins (red branches), haloarchaea globins, bacterial truncated hemoglobins (trHbO), and archaeal protoglobins. Bootstrap values are shown for some key nodes.

The gene context of *Kariarchaeaceae* globin was similar to the operon organization of *pdxS* and *pdxT* genes in some aerobic bacteria, such as *Bacillus anthracis*, *Bacillus subtilis*, and *Aquisalibacillus elongatus*. This observation prompted us to investigate whether the evolution of these operons involved horizontal gene transfer between Asgard archaea and bacteria. In the phylogenetic trees of the Pdx proteins, Asgard PdxT and PdxS both belonged to strongly supported clades together with homologs from other archaea, confidently separated from the bacterial counterparts (Fig. 2b). Thus, phylogenetic analysis indicates that Asgard PdxT and PdxS are native archaeal enzymes required for PLP biosynthesis. We then examined the evolutionary relationships among the Asgard globins, archaeal protoglobins, Haloarchaeal globin homologs (e.g. NCBI accession number: ADE04520.1) identified in this study, bacterial trHbO, and vertebrate globins. In the phylogenetic tree, all the Asgard globins and trHbO of anaerobic bacteria *Chloroflexales* formed a strongly supported clade, which was lodged within the larger bacterial trHbO branch. Archaeal protoglobins and Haloarchaeal globins formed two distinct clades, well separated from the bacterial branch with the embedded Asgard globins.

Collectively, these findings indicate that Asgard globin is a typical trHbO, likely acquired via HGT from *Chloroflexales*. Functional integration of globins with the Dxp-independent PLP biosynthesis pathway in Asgard archaea could represent an adaptive mechanism to cope with fluctuating oxygen levels in their environment.

### Enhancement of bacterial aerobic growth by Asgard globin dependent on terminal oxidase

To further characterize Asgard globins, we examined in detail the alignment of these proteins with bacterial trHbO and Haloarchaeal globins and found that 8 alpha helices with hallmarks glycine motifs, as well as heme and ligand coordinating residues ^48^, are conserved in all these proteins (Figure S2). Using AlphaFold3, we predicted structures of Asgard globin (*Kariarchaeaceae*, As_140-p_00992) and *Haloferax volcanii* globin (NCBI accession number: ADE04520.1), and found that they adopt typical 2-over-2 alpha-helical sandwich fold, which closely resembles the *Chloroflexus aurantiacus* trHbO structure (RSMD: 0.565 Å and 1.088 Å, respectively; Figure 2a). All these structures contain highly conserved heme pockets, and in particular, the proximal histidine residue that is critical for maintaining the reduced state (ferric heme) of hemoglobin ^49^ (Kari His^79^ corresponding to *H. Volcanii* His^72^ and *C. aurantiacus* His^75^). Mutation of the proximal histidine residue to phenylalanine is expected to induce oxidized state (ferrous heme), with low affinity for oxygen.

To verify the functionality of Asgard globins, the coding sequences of the *Kariarchaeaceae* archaeal globin (Kari globin for short) and its H79F mutant were codon optimized, synthesized, and inserted into a pCold-II vector. The resulting vectors were transformed into *E. coli* BL21, which was further induced for protein expression. The globins were purified and examined by optical absorption spectra (Figure 3b). The H79F mutation shifted the absorbance peak of Kari globin from 416 nm to 414 nm, consistent with the expected heme state alteration ^50^. This finding indicates that Asgard and Haloarchaeal globins possess heme pockets that are functionally analogous to the bacterial trHbO.

**Figure 3.**
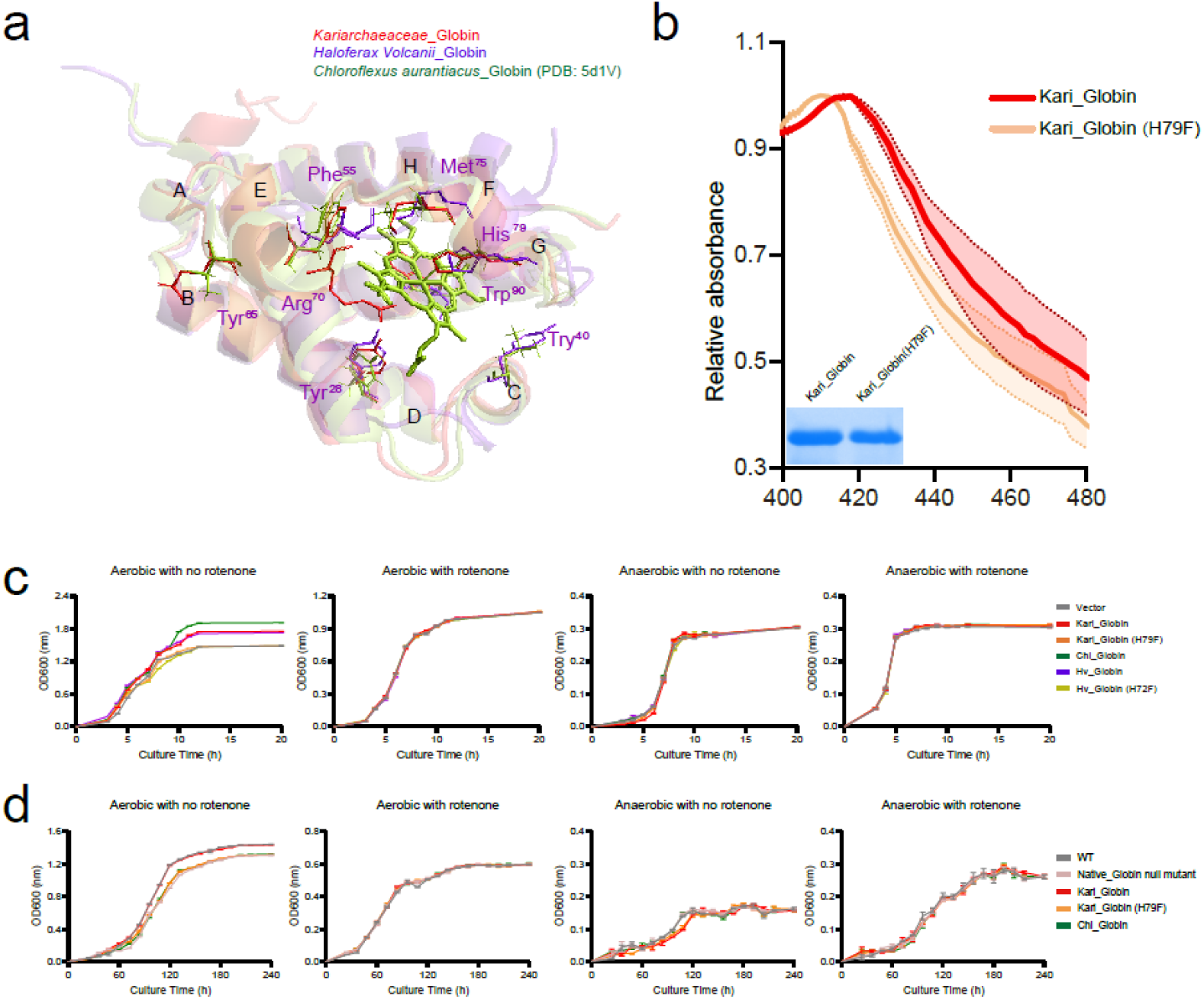
Enhancement of aerobic growth by Asgard and Haloarchaeal globins. (a) Structural comparison of Asgard globin (red), Haloarchaea globin (purple), and bacterial trHbO (green) predicted by AlphaFold3. Helical regions are labeled A-H, and residues in the highly conserved heme pockets are annotated adjacent to their positions. (b) Heme assay of wild-type *Kariarchaeaceae* globin and its H79F mutant. The lower-left inset shows SDS-PAGE analysis of purified *Kariarchaeaceae* globin and H79F mutant proteins. (c-d) Growth curves of *E. coli* (c) and *Haloferax volcanii* H1424 (d) under aerobic and anaerobic conditions, with or without rotenone treatment.

To functionally characterize the Asgard globin *in vivo*, we transformed the Kari globin and H79F mutant plasmids into *E. coli* BL21 and examined the growth of the hosts by spectrophotometry under aerobic and anaerobic conditions. In addition to the Asgard globin, the *H. volcanii* globin and its oxidized-mutant (H72F), as well as *C. aurantiacus* globin were included in the assay. To perform the growth assay, *E. coli* was first aerobically cultured (ambient atmosphere, 180 rpm shaking, 37 °C) and then inoculated into fresh medium at a 1:1000 dilution ratio before further growth assays. The growth experiments show that expression of Kari globin, but not the H79F mutant, significantly enhanced growth of *E. coli* under aerobic condition (one-way ANOVA analysis, *p* <0.01), whereas no difference in growth was observed in the absence of oxygen (Figure 3c). These findings prompted us to explore whether these globin proteins enhanced aerobic growth through terminal oxidase. To address this question, we performed additional growth assays after adding Rotenone (10 μM). Rotenone is a respiration inhibitor that blocks electron transfer to terminal oxidase ^51^. The results showed that, under aerobic conditions, rotenone inhibited *E. coli* growth and completely abolished the effect of Asgard globin, indicating that enhancement of *E. coli* growth by Asgard globin depended on the terminal oxidase analogously to their bacterial counterparts, trHbO.

To examine the biological function of Asgard globins in an archaeal cellular context, we genetically replaced the native globin gene of *H. volcanii* H1424 with Asgard globin and its H79F mutant, as well as *C. aurantiacus* globin. As a control, we generated a globin null mutant by deleting the native globin gene in *H. volcanii*. The growth curves of these *H. volcanii* mutants were determined at 42 °C (optimal growth temperature of *H. volcanii*) under both aerobic and anaerobic conditions. The spectrophotometry results showed that the native globin null mutation caused a significant aerobic growth defect of *H. volcanii* (one-way ANOVA analysis, *p*< 0.01; Figure 3d). The aerobic growth was restored by Kari globin, whereas no growth recovery occurred with the Kari H79F mutant or *C. aurantiacus* globin. The growth difference between the wildtype *H. volcanii* and its mutants was eliminated by Rotenone addition (10 μM) or under anaerobic conditions, consistent with the observations in *E. coli*. Thus, Asgard globins contribute to aerobic growth of archaea via terminal oxidases.

Overall, the results of the biochemical, and genetic analyses demonstrate that the Asgard and Haloarchaeal globins enhance aerobic growth dependent on terminal oxidase. The functional plasticity of the globins across bacteria and archaea likely enables efficient horizontal gene transfer and thus facilitates rapid host adaptation to oxygen dynamics.

### Switching expression of Asgard globin-related promoter region between aerobic and anaerobic conditions

Considering the role of Asgard and *H. volcanii* globins in aerobic growth, we wondered whether the promoter regions of these globins responded to environmental oxygen. To address this question, we first characterized the promoter region of the native globin in *H. volcanii* under aerobic and anaerobic conditions using reverse transcription quantitative PCR (RT-qPCR). *H. volcanii* was grown in normal atmospheric oxygen condition for aerobic expression analysis. The RT-qPCR data showed that the Haloarchaeal globin expression was strongly driven by aerobic condition, whereas under anaerobic conditions, its transcript level decreased by about 20- to 200-fold (Figure 4a). This oxygen-responsive promoter activity of the Haloarchaeal globin gene is consistent with its role in supporting aerobic growth.

**Figure 4.**
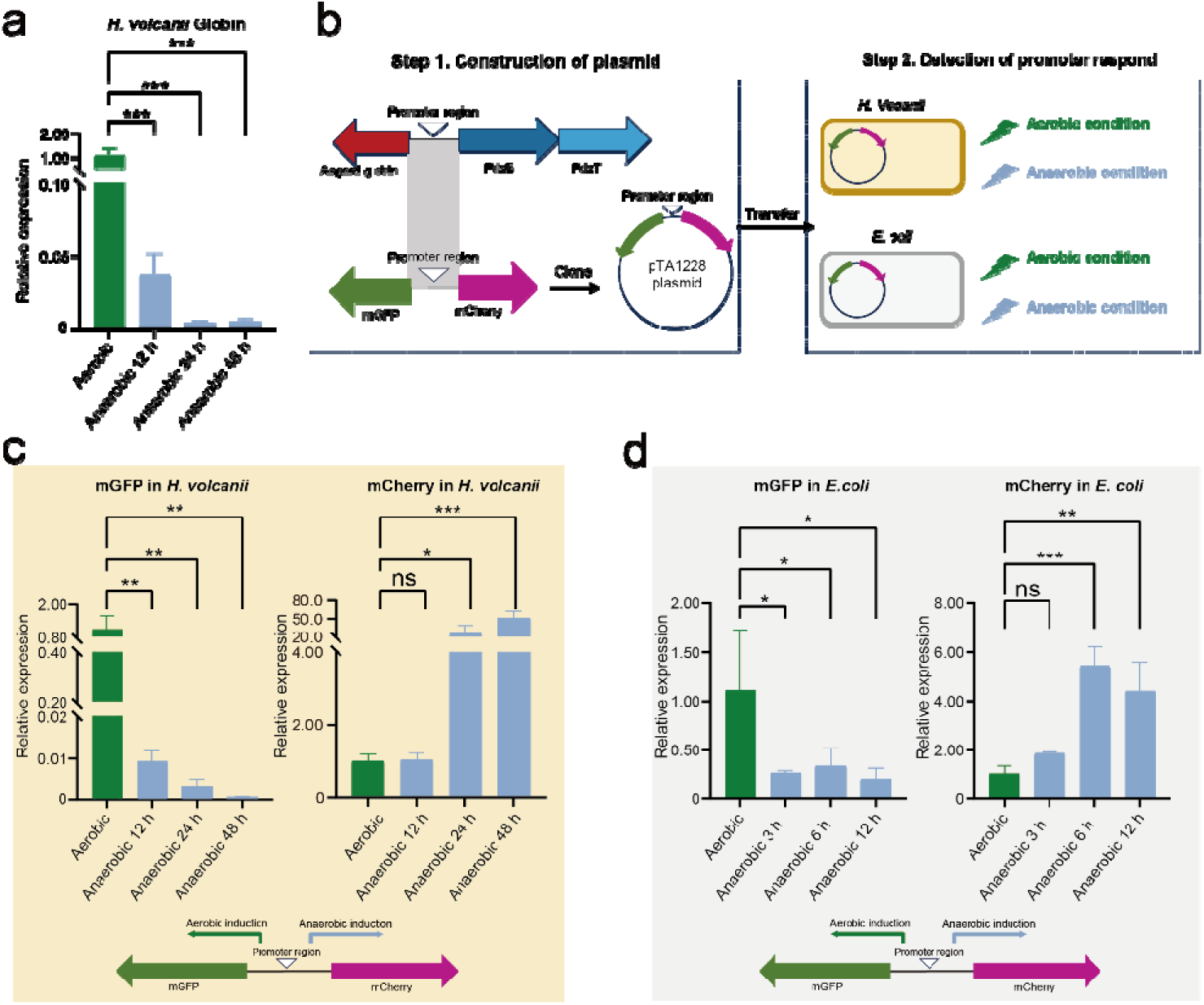
Oxygen-induced expression of Asgard and Haloarchaeal globins. (a) Endogenous globin mRNA expression levels in *Haloferax volcanii* H1424 under aerobic (green bars) and anaerobic (blue bars) conditions. The y-axis is truncated to visualize the extremely low relative expression of globin under anaerobic conditions. Asterisks indicate significant differences: *p<0.05, **p<0.01, and, ***p<0.001; ns denotes non-significant differences (same for all following figures). (b) Experimental flowchart for analyzing Asgard globin expression under different oxygen conditions in *Haloferax volcanii* H1424 (*H. volcanii*) (c), and *Escherichia coli* (*E. coli*), (c-d) Expression of the Asgard globin promoter region in *H. volcanii* (c) and *E. coli* (d) under aerobic and anaerobic conditions.

To further characterize the Asgard globin-related promoter region and rule out potential effects of its archaeal protein products on the heterogeneous cellular system, we synthesized the native promoter region between the Asgard globin and PdxS genes from *Kariarchaeaceae*, and inserted this sequence into a dual-fluorescence reporter system by fusing mGFP and mCherry coding sequences to its 5’ and 3’ termini, respectively (Figure 4b). Then, this reporter cassette was cloned into a shuttle vector pTA1228, and transformed into either *H. volcanii* or *E. coli* BL21 for subsequent RT-qPCR analysis under aerobic and anaerobic conditions. In *H. volcanii* cell, the mGFP showed pronounced aerobic induction, with expression levels >5-fold higher under aerobic versus anaerobic conditions (Figure 4c). While mCherry expression remained low under both aerobic and anaerobic conditions at 12 h, it exhibited strong anaerobic induction at 24 h and 48 h, with a ∼200-fold increase in expression. Collectively, these results demonstrate that the Asgard globin is aerobically induced through activation of its promoter that functions as an oxygen-sensing molecular switch in Asgard archaea.

Given the apparent bacterial origin of Asgard globins and their potential role in enhancing bacterial aerobic growth, we also assessed the activity of its promoter region in *E. coli* using the above reporter cassette. The RT-qPCR analysis showed that, under aerobic conditions, mGFP expression increased about 4-fold (3 h), 3-fold (6 h), and 6-fold (12 h), compared to anaerobic conditions. In contrast, mCherry expression was anaerobically induced, increasing by about 5-fold increase at 6 h and 12 h, relative to that under aerobic conditions. These findings demonstrate functional robustness of the Asgard globin promoter region for sensing the environment oxygen level in both archaea and bacteria.

Taken together, these findings indicate that the promoters of Haloarchaea globin and Asgard globin can be aerobically activated, compatible with the role of the globins in promoting aerobic growth. Under anaerobic conditions, the promoter region of the Asgard globin gene downregulates globin expression, while inducing PdxS and PdxT expression required for PLP biosynthesis. This sophisticated regulatory mechanism likely facilitates Asgard adaptation to both aerobic and anaerobic environments. The functionality of the Asgard globin promoter region in *E. coli* is compatible with the bacterial origin of this gene as suggested by phylogenetic analysis (Figure 2c).

## DISCUSSION

This study reveals the critical roles of terminal oxidase and globin in oxygen-adaptive flexibility of Asgard archaea. Although there is no indication that Asgard archaea thrive in oxic niches, trace oxygen could induce their globin expression, and thereby promote growth under nanoxic condition, by stimulating terminal oxidase. Conversely, strict anaerobic conditions shift Asgard metabolic states, in particular, by downregulating globin expression and activating PLP biosynthesis pathway to compensate for the lack of oxygen.

Genome comparison, structural modelling, and phylogenetic analyses complemented by biochemical assays show that certain Asgard lineages, including *Kariarchaeaceae*, *Gerdarchaeales*, and *Hodarchaeales*, encodes active A-type terminal oxidases. The monophyly of Asgard CyoB, along with its counterparts in TACK archaea and Haloarchaea, suggests a long history of vertical evolution of terminal oxidases in archaea and in Asgard in particular. Thus, Asgard archaea including *Hodarchaeales* that are currently considered to be the closest known relatives of eukaryotes apparently adapted to oxygen at an early stage of their evolution.

In contrast to the Haloarchaeal globin that forms a distinct clade in the phylogenetic tree, well separated from other clades, phylogenetic analysis and protein structure comparison strongly suggest the origin of the globins of *Kariarchaeaceae* and *Hodarchaeales* by horizontal transfer from *Chloroflexi* bacteria. Consistently, previous studies reported transfer of multiple genes between archaea and *Chloroflexi* ^52,53^. The *Chloroflexales* are typically anaerobic, phototrophic or chemoheterotrophic bacteria that inhabit mesothermal freshwater or moderate hot spring environments ^54^ where they may coexist with Asgard archaea. Combined with the previous observation that *Archaeplastida* (plant and algal) hemoglobin apparently evolved from within *Chloroflexales* ^56,57^, our findings are compatible with the possibility imply that horizontal transfer of the bacterial globin gene into Asgard archaea predated the origin of eukaryotes.

The results of this work point to oxygen tolerance of some Asgard archaea and elaborate regulation of the expression of the relevant genes during aerobic and anaerobic growth. Under aerobic conditions, the Asgard globin promoter shows strongly enhanced activity, promoting growth supported by the terminal oxidase-dependent pathway. Similar oxygen activation of the globin promoter was observed in this work for Haloarchaea and previously for *Mycobacterium tuberculosis* ^20^. Given that oxygen is required for the activity of some PLP-dependent enzymes, anaerobic upregulation of Asgard PdxS and PdxT could represent a compensatory mechanism to sustain the homeostasis of PLP-dependent proteins under oxygen-limited conditions. The functional robustness of globin, terminal oxidase, the PLP pathway, and their regulatory mechanisms might reflect evolution of Asgard archaea under conditions of fluctuating environmental oxygen levels ^58^.

The key finding of this work, the oxygen-adaptive plasticity of Asgard archaea, is a trait that was likely crucial for eukaryogenesis. Although genomic reconstruction and cultivation studies suggest an anaerobic common ancestor for Asgard archaea^33,37,59,60^, our present findings point to the possibility that the Asgard lineage ancestral to eukaryotes evolved oxygen tolerance. This lineage likely exploited nanomolar oxygen levels analogously to nanaerobic microorganisms via Uox terminal oxidases, possibly, acquired from other archaea (Figure 5). These terminal oxidases could reduce oxygen by utilizing electrons generated by distinct functional units, such as microcompartments (Du et al. 2025). Nanoxic conditions resemble the weakly oxygenated Proterozoic oxygen minimum zones, where eukaryogenesis is thought to have occurred ^4,63^. The apparent acquisition of bacterial globin by Asgard archaea emphasizes the contribution of early bacterial HGT to eukaryogenesis. Adaptation of Asgard archaea to oxygen influx in nanoxic niches likely facilitated the initial endosymbiotic interaction between the Asgard ancestor of eukaryotes and the alphaproteobacterial proto-mitochondrion. During eukaryogenesis, the Asgard Uox apparently was displaced by the alphaproteobacterial terminal oxidases that became the mitochondrial Cox.

**Figure 5.**
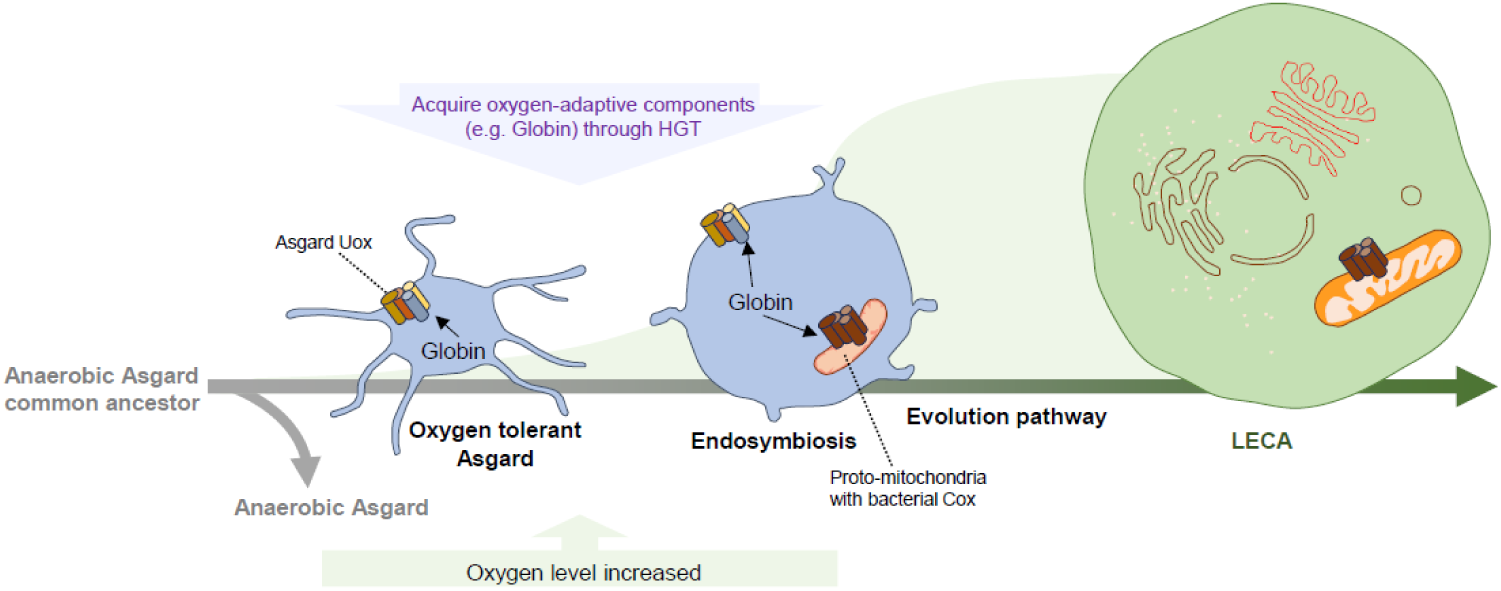
Proposed model of oxygen-driven eukaryogenesis. In this model, the Asgard common ancestor had a strictly anaerobic lifestyle. Subsequently, adapting for survival in the increasingly oxygenated environment, some Asgard lineages evolved tolerance to trace amounts of oxygen through the acquisition of a Uox-family terminal oxidase, likely, from other archaea and oxygen-adaptive components such as globin, from bacteria. The Asgard globin might have activated alphaproteobacterial proto-mitochondrial Cox, facilitating endosymbiosis and eukaryogenesis. After endosymbiosis, the Asgard Uox was replaced by the proto-mitochondrial Cox antedating the emergence of the last eukaryotic common ancestor (LECA).

In conclusion, this work presents evidence that some lineages of Asgard archaea including *Hodarchaeales*, the apparent closest known relatives of eukaryotes, evolved oxygen tolerance conferred by a terminal oxidase system, likely aided by globins. Our findings suggest that eukaryogenesis initially occurred under nanoxic conditions where the Asgard archaeal host and the alphaproteobacterial ancestors of the mitochondria could both thrive and productively interact. Future research combining advanced cell biology with cultivation efforts will be essential to elucidate the ecological and evolutionary plasticity of Asgard archaea and their contributions to eukaryotic origin.

## Acknowledgments

*Haloferax volcanii* H1424 and pTA1228 vector was kindly provided by Hua Xiang (Institute of Microbiology, Chinese Academy of Sciences Institute of Microbiology, Chinese Academy of Sciences). This work was supported by National Natural Science Foundation of China (No.32225003, 32393970, 32393971, 32370004, 32393973, 92051102, and 92251306), Guangdong Basic and Applied Basic Research Foundation (No. 2023A1515011309), Shenzhen Medical Research Fund (No. B2301005), Guangdong Major Project of Basic and Applied Basic Research (No. 2023B0303000017), Project of Department of Education of Guangdong Province (No. 2025KCXTD039), Shenzhen Natural Science Fund (No. 20220809161641002), Shenzhen University 2035 Program for Excellent Research (No. 2022B002), the Synthetic Biology Research Center of Shenzhen University and Scientific Foundation for Youth Scholars of Shenzhen University.

## Author contributions

Z.L., R.X., and M.L. conceived and designed the experiments. R.X., A.X., and H.C. performed the experiments. Z.L., R.X., and Y.L. performed the bioinformatics analyses. Z.L., and R.X. analyzed the data. Z.L., R.X., and M.L. wrote the paper, and all authors edited and approved the paper.

## METHODS

### Bioinformatics analysis

All the sequences were obtained by our Asgard database ^37^ or National Center for Biotechnology Information database. The protein sequences were aligned using MUSCLE (Ver 3.8.1551) ^64^, trimmed with TrimAI (Ver 1.4) before construction of phylogenetic tree using IQ-tree (Ver 1.6.12) ^65^. The three-dimensional structures for Asgard globin and CyoA, CyoB, CyoC, and CyoD were built by AlphaFold3 ^45^.

### Protein expression and purification

The coding sequences of *Kariarchaeaceae* globin, *Kariarchaeaceae* globin mutant (H79F), Haloarchaeal globin, Haloarchaeal globin mutant (H72F), and *Chloroflexus aurantiacus* TrHbO (NCBI accession number: WP_012256033.1) were codon optimized for expression in *E. coli*, synthesized, and respectively, assembled into a pCold-II vector (TaKaRa, Japan) that contains an N-terminal His tag. The recombinant vectors were respectively transformed into *E. coli* BL21 strains before inducible expression. Briefly, the BL21 strain carrying the recombinant vector was incubated in Luria Broth medium at 37 °C until the optical density OD_600_ reached about 0.6, and then isopropyl-D-1-thiogalactopyranodside and was added at a final concentration of 0.1 mM, additionally, 0.02 g/L FeCl₃ was added to facilitate heme synthesis., followed by incubation at 15 °C for about 18 hours. Then, the target proteins were purified by ÄKTA Pure System (GE Healthcare, USA) with Ni-NTA affinity chromatography column (HisTrap HP, GE Healthcare, USA). Next, the purified proteins were pooled and concentrated in 20 mM phosphate buffer solution (pH 7.4) by Amicon Ultra-15 (Millipore, USA). The SDS-PAGE was used to detect the targeted proteins.

### Terminal oxidase activity Assay

To examine the activity of the Hodarchaeales_NJBF01000076.1 terminal oxidase, genes *cyoA* and *cyoD* were codon-optimized and cloned into the pCDF-Duet vector, while *cyoB* (or its H169F mutant gene) and *cyoC* were similarly codon-optimized and inserted into the pRSF-Duet vector. The recombinant vectors were co-transformed into *E. coli* BL21 (DE3) strains and incubated in LB medium at 37 °C. Once the optical density reached ∼0.6, 0.5 mM IPTG and 0.25 mM CuSO₄ were added, followed by incubation at 37 °C for 3–4 hours. Cells were centrifuged at 5,000 rpm for 15 minutes and resuspended in 20 mL of 50 mM K-phosphate buffer (pH 7.5). The resuspended cells were sonicated at 180 W for 25 minutes using intermittent cycles (2 seconds on, 3 seconds off). The lysate was then centrifuged at 10,000 g and 4°C for 20 minutes to remove debris. The supernatant was transferred to ultracentrifuge tubes and subjected to high-speed centrifugation at 100,000 g for 2 hours. The resulting pellet was dissolved in the same buffer supplemented with 2% Triton X-100 and incubated at 4°C with shaking for 2 hours to solubilize membrane proteins.

The cytochrome c oxidase activity measurement was done according to the instruction manual of Cytochrome C Oxidase Assay Kit (Abcam, ab239711, UK). In short, cytochrome c was reduced with DTT for 15 minutes, and the absorbance at 550 nm was verified to fall within 0.2–0.6. Subsequently, 10 μL solubilized membrane proteins (empty vector control, experimental group, or mutant group) were mixed with 120 μL reduced cytochrome c, and absorbance at 550 nm was immediately monitored every 10 seconds for 35 minutes using a multifunctional spectrophotometer.

Quinol oxidase activity was assessed by measuring absorbance changes at 275 nm as previously described ^66^, in a reaction mixture containing 50 mM sodium phosphate buffer (pH 7.5), 30 μM antimycin A, and 75 μM reduced quinol ^67^. Specific activity was calculated using the molar extinction coefficient of cytochrome c (7.04 mM^-1^cm^-1^)/ quinone (0.4 mM⁻¹ cm⁻¹) and total protein concentration determined by BCA assay (Thermo Fisher Scientific, USA).

### Heme assay

The heme experiments were conducted according to previous method ^68^. Culture *E. coli* BL21 strains expressing Asgardglobin and mutant proteins (H79F) separately for protein overexpression. Centrifuge to collect the induced bacterial cells (0.1 g wet weight of pellet), and wash twice with minimal salt medium (60 mM K₂HPO₄, 33 mM KH₂PO₄, 7.6 mM (NH₄)₂SO₄, and 1.7 mM sodium citrate). Resuspend the pellet in 0.6 mL alkaline pyridine reagent (2.2 M pyridine, 0.1 M NaOH) and sonicate to disrupt the cells. Centrifuge the lysate at 25,000 g for 10 minutes at 4 °C, and collect the supernatant. Scan the absorption spectrum of the supernatant between 360–600 nm to detect characteristic peaks corresponding to the reduced state (Fe²⁺, Ferrous-trHbO) and oxidized state (Fe³⁺, Ferri-trHbO) of heme-bound iron.

### Measurment of growth curve

*E. coli* BL21 cells bearing pCold-II (empty vector) and Asgard globin, Asgard globin (H79F), *Chloroflexus aurantiacus* TrHbO, *H. volcanii* globin, and *H. volcanii* globin (H72F) were cultured in LB medium overnight at 37 °C, 180 rpm. On the second day, inoculate at a 1:1000 ratio into LB medium containing antibiotics and 0.02 g/L FeCl₃ in a glass tube. For anaerobic culture conditions, supplement with an additional 20 mM KNO₃ (electrons acceptor). The anaerobic cultivation vessel, which was 27 mL in volume and contained 15 mL medium, was sealed with butyl rubber; the gas phase in the vessel was then exchanged by gentle bubbling with N_2_ gas for 15 minutes using a sterile needle to remove dissolved oxygenbefore sterilization. The cultivation vessels were shaken at 80 rpm at 37 °C in the dark. Rotenone was dissolved in DMSO to a final concentration of 1 mM, and further diluted 100-fold to achieve a working concentration of 10 µM.

To measure the growth curve of Asgard globin within archaeal cellular environment, we selected the model halophilic archaeon *Haloferax volcanii* H1424. *Haloferax volcanii* H1424 and its derived strains *Δ*globin (native globin knockout), Asgard-globin (native globin replaced by Asgard globin), H79F-globin (native globin replaced by Asgard globin H79F mutation), and Bacterial-globin (native globin replaced by *Chloroflexus aurantiacus* TrHbO) were cultured aerobically in Hv-YPC medium at 42 °C, 180 rpm overnight. Cultures were subcultured (1:1000 dilution) into fresh medium for growth curve measurements. For anaerobic conditions, cells were cultivated following *E. coli* anaerobic protocols, except supplemented with 50 mM KNO₃. All experiments were performed with three biological replicaed per group.

### mRNA extraction and reverse transcription quantitative PCR

The native promoter region between the Asgard globin and PdxS genes in *Kariarchaeaceae* genomes flanked by mGFP and mCherry coding sequences at its 5’ termini and 3’ termini was synthesized and cloned into the shuttle vector pTA1228, generating the plasmid designated pTA1228-GloPro. *E. coli* BL21 and *H. volcanii* H1424 cells harboring the pTA1228-GloPro plasmid were cultured in 20 mL of medium to the logarithmic phase (OD_600_ = 0.6–0.8). The H1424 culture was divided into 10 mL aliquots in sterile anaerobic culture tubes, supplemented with 50 mM KNO_3_ (for *E.coli* it was 20 mM KNO_3_), and flushed with nitrogen gas budding for 10 minutes using a sterile needle. Cultures were further incubated at 37 °C with gentle shaking (80 rpm) for 12 hours, 24 hours, and 48 hours (for *E. coli* it was 3 hours, 6 hours, and 12 hours). Cells were harvested by centrifugation at 5,000 rpm for 5 minutes at 4 °C, and total RNA was extracted using Direct-zol^TM^ RNA Miniprep (Zymo research, USA). The H1424 cell lysis was achieved by vigorous vortexing with sterilized zirconia grinding beads (0.1 mm diameter) for 10 minutes. Following centrifugation at 13,000 × g at 4°C for 10 minutes, the supernatant was collected for total RNA extraction using the Direct-zol™ RNA MiniPrep kit (ZYMO research, USA).

Extracted RNA was diluted to a concentration of 100 ng/μL. Reverse transcription and quantitative PCR (RT-qPCR) were performed in 20 μL reaction volumes using the UniPeak U+ One Step RT-qPCR SYBR Green Kit (Vazyme, China). Amplification and fluorescence detection were carried out on a QuantStudio 3 Real-Time PCR System (Thermo Fisher Scientific, USA). The relative mRNA expression levels under aerobic and anaerobic conditions were calculated using the 2^−ΔΔCt^ method. All experiments were conducted with three biological replicates.

### Genetic manipulation of *Haloferax volcanii*

To construct *H. volcanii* mutants, the upstream and downstream homologous arms of the *globin* gene in *H. volcanii* H1424 were amplified respectively, with H1424 genomic DNA as the template. The *pyrE2* selectable marker gene was amplified from the pTA1228 vector. For gene replacement, the target gene was inserted between the upstream homologous arm and the *pyrE2* marker. The replaced coding region was codon-optimized, synthesized, and assembled into the pTA1228 vector. A linear targeting cassette was constructed via overlap PCR.

Transformation of the resulted cassettes to *H. volcanii* H1424 was carried out according to a previously described protocol ^69^. Briefly, 10 mL *Haloferax volcanii* H1424 was grown in Hv-YPC medium to an OD₆₀₀ of ∼0.8. Cells were pelleted by centrifugation at 8,000 rpm and washed once with buffered spheroplasting solution. Resuspended the pellet in 600 μL buffered protoplasting solution and aliquot into three tubes (200 μL per tube). Gently added 20 μL 0.5 M EDTA, mix by horizontal agitation, and incubated at room temperature (RT) for 10 minutes to allow spheroplast formation. Added 30 μL DNA mixture (containing 1–2 μg DNA, 15 μL unbuffered spheroplasting solution, and 5 μL 0.5 M EDTA) to each tube. Added 250 μL 60% PEG solution to each tube. Mixed gently by horizontal shaking for 10 minutes, then incubated at RT for 30 minutes. Added 1.5 mL spheroplast dilution solution, mixed, and incubated at RT for 2 minutes. Centrifuged at 8,000 rpm and discarded the supernatant. Resuspended the pellet in 1 mL cell recovery solution and incubated at 42 °C for 1.5–2 hours. Resuspended by pipetting and incubated at 42 °C with shaking at 180 rpm for 3–4 hours. Centrifuged at 8,000 rpm and washed twice with transformation dilution solution to eliminate uracil. Resuspended cells in 1 mL transformation dilution solution, prepared 10-fold serial dilutions, and plated 100 μL onto Hv-Ca medium. Transformants were incubated and single colonies were selected for validation by colony PCR and DNA sequencing.

---

**Figure S1.**
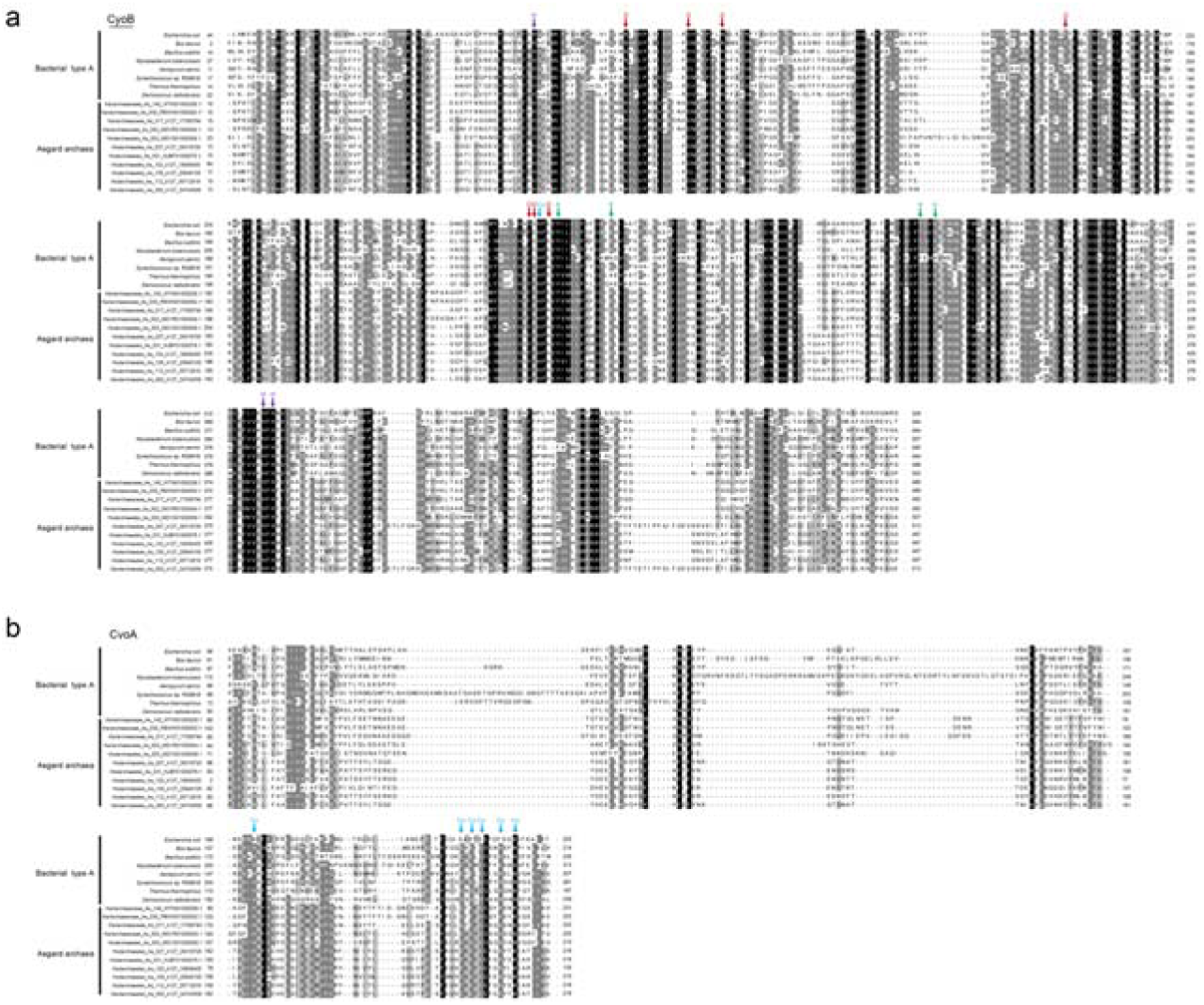
Amino acid sequence alignment of CyoB (a) and CyoA (b) in Asgard archaea and bacteria. The D- and K- proton channels, and histidine ligands to heme and Cu_B_ in CyoB, and Cu_A_-binding histidine ligands in CyoA are labelled.

**Figure S2.**
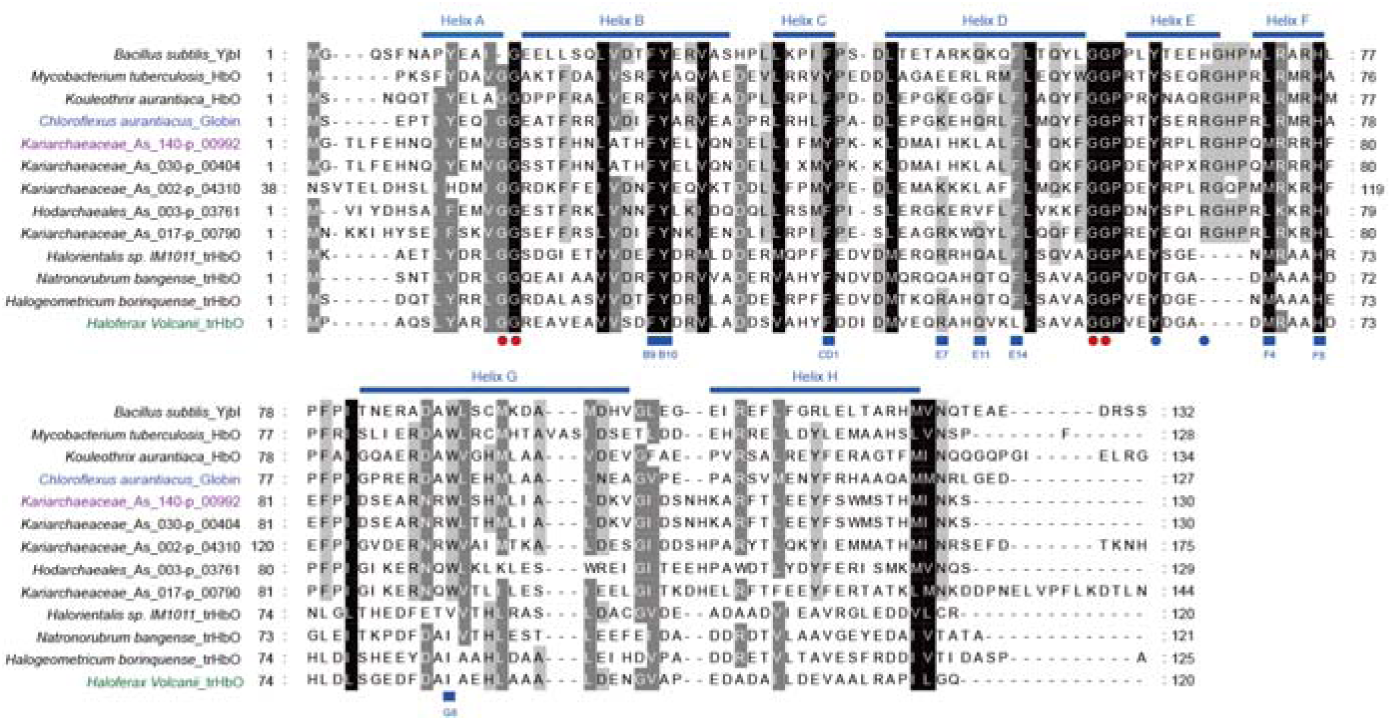
Sequence alignment of Asgard globins, Haloarchaeal globins, and bacterial trHbO. The predicted conserved helical regions are indicated by blue rectangles above the aligned sequences, with glycine motifs (blue circles), heme-coordinating residues (red circles), and ligand-binding residues (blue rectangles) highlighted below the aligned sequences.

## References

1 Gross, J. & Bhattacharya, D. Uniting sex and eukaryote origins in an emerging oxygenic world. Biology Direct 5, 1–20 (2010).

2 Luo, G. et al. Rapid oxygenation of Earth’s atmosphere 2.33 billion years ago. Sci Adv 2, e1600134, doi:10.1126/sciadv.1600134 (2016).

3 Knoll, A. H., Bergmann, K. D. & Strauss, J. V. Life: the first two billion years. Philos Trans R Soc Lond B Biol Sci 371, doi:10.1098/rstb.2015.0493 (2016).

4 Mills, D. B. et al. Eukaryogenesis and oxygen in Earth history. Nature ecology & evolution 6, 520–532 (2022).

5 Lane, N. & Martin, W. The energetics of genome complexity. Nature 467, 929–934 (2010).

6 Booth, A. & Doolittle, W. F. Eukaryogenesis, how special really? Proceedings of the National Academy of Sciences 112, 10278–10285 (2015).

7 Lu, Z. & Imlay, J. A. When anaerobes encounter oxygen: mechanisms of oxygen toxicity, tolerance and defence. Nat Rev Microbiol 19, 774–785, doi:10.1038/s41579-021-00583-y (2021).

8 Khademian, M. & Imlay, J. A. How Microbes Evolved to Tolerate Oxygen. Trends Microbiol 29, 428–440, doi:10.1016/j.tim.2020.10.001 (2021).

9 Sousa, F. L. et al. The superfamily of heme-copper oxygen reductases: types and evolutionary considerations. Biochim Biophys Acta 1817, 629–637, doi:10.1016/j.bbabio.2011.09.020 (2012).

10 Pereira, M. M., Santana, M. & Teixeira, M. A novel scenario for the evolution of haem-copper oxygen reductases. Biochim Biophys Acta 1505, 185–208, doi:10.1016/s0005-2728(01)00169-4 (2001).

11 Esposti, M. D. On the evolution of cytochrome oxidases consuming oxygen. Biochim Biophys Acta Bioenerg 1861, 148304, doi:10.1016/j.bbabio.2020.148304 (2020).

12 Gao, Y. et al. Heme-copper terminal oxidase using both cytochrome c and ubiquinol as electron donors. Proceedings of the National Academy of Sciences 109, 3275–3280 (2012).

13 Dikshit, K. L., Spaulding, D., Braun, A. & Webster, D. A. Oxygen inhibition of globin gene transcription and bacterial haemoglobin synthesis in Vitreoscilla. Microbiology 135, 2601–2609 (1989).

14 Pawaria, S., Lama, A., Raje, M. & Dikshit, K. L. Responses of Mycobacterium tuberculosis hemoglobin promoters to in vitro and in vivo growth conditions. Applied and environmental microbiology 74, 3512–3522 (2008).

15 Miralles-Robledillo, J. M., Martinez-Espinosa, R. M. & Pire, C. Transcriptomic profiling of haloarchaeal denitrification through RNA-Seq analysis. Appl Environ Microbiol 90, e0057124, doi:10.1128/aem.00571-24 (2024).

16 Liu, C., He, Y. & Chang, Z. Truncated hemoglobin o of Mycobacterium tuberculosis: the oligomeric state change and the interaction with membrane components. Biochemical and biophysical research communications 316, 1163–1172 (2004).

17 Hardison, R. C. A brief history of hemoglobins: plant, animal, protist, and bacteria. Proceedings of the National Academy of Sciences of the United States of America 93, 5675 (1996).

18 Milani, M., Pesce, A., Ouellet, H., Guertin, M. & Bolognesi, M. Truncated hemoglobins and nitric oxide action. IUBMB Life 55, 623–627, doi:10.1080/15216540310001628708 (2003).

19 Freitas, T. A. K., Saito, J. A., Hou, S. & Alam, M. Globin-coupled sensors, protoglobins, and the last universal common ancestor. Journal of inorganic biochemistry 99, 23–33 (2005).

20 Pawaria, S., Lama, A., Raje, M. & Dikshit, K. L. Responses of Mycobacterium tuberculosis hemoglobin promoters to in vitro and in vivo growth conditions. Appl Environ Microbiol 74, 3512–3522, doi:10.1128/AEM.02663-07 (2008).

21 Yu, F. et al. Recent advances in the physicochemical properties and biotechnological application of Vitreoscilla hemoglobin. Microorganisms 9, 1455 (2021).

22 Hoffarth, E. R., Rothchild, K. W. & Ryan, K. S. Emergence of oxygen- and pyridoxal phosphate-dependent reactions. FEBS J 287, 1403–1428, doi:10.1111/febs.15277 (2020).

23 Bisello, G., Longo, C., Rossignoli, G., Phillips, R. S. & Bertoldi, M. Oxygen reactivity with pyridoxal 5’-phosphate enzymes: biochemical implications and functional relevance. Amino Acids 52, 1089–1105, doi:10.1007/s00726-020-02885-6 (2020).

24 Fitzpatrick, T. B. et al. Two independent routes of de novo vitamin B6 biosynthesis: not that different after all. Biochem J 407, 1–13, doi:10.1042/BJ20070765 (2007).

25 Mittenhuber, G. Phylogenetic analyses and comparative genomics of vitamin B6 (pyridoxine) and pyridoxal phosphate biosynthesis pathways. J Mol Microbiol Biotechnol 3, 1–20 (2001).

26 Bertagnolli, A. D. & Stewart, F. J. Microbial niches in marine oxygen minimum zones. Nat Rev Microbiol 16, 723–729, doi:10.1038/s41579-018-0087-z (2018).

27 Berg, J. S. et al. How low can they go? Aerobic respiration by microorganisms under apparent anoxia. FEMS Microbiology Reviews 46, fuac006 (2022).

28 Baughn, A. D. & Malamy, M. H. The strict anaerobe Bacteroides fragilis grows in and benefits from nanomolar concentrations of oxygen. Nature 427, 441–444, doi:10.1038/nature02285 (2004).

29 Lee, S. H., Youn, H., Kang, S. G. & Lee, H. S. Oxygen-mediated growth enhancement of an obligate anaerobic archaeon Thermococcus onnurineus NA1. J Microbiol 57, 138–142, doi:10.1007/s12275-019-8592-y (2019).

30 Vosseberg, J. et al. The emerging view on the origin and early evolution of eukaryotic cells. Nature 633, 295–305, doi:10.1038/s41586-024-07677-6 (2024).

31 Lopez-Garcia, P. & Moreira, D. The Syntrophy hypothesis for the origin of eukaryotes revisited. Nat Microbiol 5, 655–667, doi:10.1038/s41564-020-0710-4 (2020).

32 Spang, A. et al. Proposal of the reverse flow model for the origin of the eukaryotic cell based on comparative analyses of Asgard archaeal metabolism. Nature microbiology 4, 1138–1148 (2019).

33 Eme, L. et al. Inference and reconstruction of the heimdallarchaeial ancestry of eukaryotes. Nature 618, 992–999, doi:10.1038/s41586-023-06186-2 (2023).

34 Spang, A. et al. Complex archaea that bridge the gap between prokaryotes and eukaryotes. Nature 521, 173–179, doi:10.1038/nature14447 (2015).

35 Zaremba-Niedzwiedzka, K. et al. Asgard archaea illuminate the origin of eukaryotic cellular complexity. Nature 541, 353–358 (2017).

36 Imachi, H. et al. Isolation of an archaeon at the prokaryote–eukaryote interface. Nature 577, 519–525 (2020).

37 Liu, Y. et al. Expanded diversity of Asgard archaea and their relationships with eukaryotes. Nature 593, 553–557, doi:10.1038/s41586-021-03494-3 (2021).

38 Eme, L. et al. Inference and reconstruction of the heimdallarchaeial ancestry of eukaryotes. Nature, 1–8 (2023).

39 Zhang, J. et al. Deep origin of eukaryotes outside Heimdallarchaeia within Asgardarchaeota. Nature, doi:10.1038/s41586-025-08955-7 (2025).

40 Seitz, K. W. et al. Asgard archaea capable of anaerobic hydrocarbon cycling. Nat Commun 10, 1822, doi:10.1038/s41467-019-09364-x (2019).

41 Bulzu, P. A. et al. Casting light on Asgardarchaeota metabolism in a sunlit microoxic niche. Nat Microbiol 4, 1129–1137, doi:10.1038/s41564-019-0404-y (2019).

42 Appler, K. E. et al. Oxygen metabolism in descendants of the archaeal-eukaryotic ancestor. BioRxiv, 2024.2007. 2004.601786 (2024).

43 Imachi, H. et al. Eukaryotes’ closest relatives are internally simple syntrophic archaea. bioRxiv, 2025.2002. 2026.640444 (2025).

44 Su, C. C. et al. A ‘Build and Retrieve’ methodology to simultaneously solve cryo-EM structures of membrane proteins. Nat Methods 18, 69–75, doi:10.1038/s41592-020-01021-2 (2021).

45 Abramson, J. et al. Accurate structure prediction of biomolecular interactions with AlphaFold 3. Nature 630, 493–500, doi:10.1038/s41586-024-07487-w (2024).

46 Abramson, J. et al. The structure of the ubiquinol oxidase from Escherichia coli and its ubiquinone binding site. Nat Struct Biol 7, 910–917, doi:10.1038/82824 (2000).

47 Kozubal, M. A., Dlakic, M., Macur, R. E. & Inskeep, W. P. Terminal oxidase diversity and function in “Metallosphaera yellowstonensis”: gene expression and protein modeling suggest mechanisms of Fe(II) oxidation in the sulfolobales. Appl Environ Microbiol 77, 1844–1853, doi:10.1128/AEM.01646-10 (2011).

48 Jamil, F. et al. Crystal structure of truncated haemoglobin from an extremely thermophilic and acidophilic bacterium. The journal of biochemistry 156, 97–106 (2014).

49 Bonaventura, C., Henkens, R., Alayash, A. I., Banerjee, S. & Crumbliss, A. L. Molecular controls of the oxygenation and redox reactions of hemoglobin. Antioxid Redox Signal 18, 2298–2313, doi:10.1089/ars.2012.4947 (2013).

50 Fabozzi, G., Ascenzi, P., Renzi, S. D. & Visca, P. Truncated hemoglobin GlbO from Mycobacterium leprae alleviates nitric oxide toxicity. Microb Pathog 40, 211–220, doi:10.1016/j.micpath.2006.01.004 (2006).

51 Fato, R. et al. Differential effects of mitochondrial Complex I inhibitors on production of reactive oxygen species. Biochim Biophys Acta 1787, 384–392, doi:10.1016/j.bbabio.2008.11.003 (2009).

52 Meheust, R., Castelle, C. J., Jaffe, A. L. & Banfield, J. F. Conserved and lineage-specific hypothetical proteins may have played a central role in the rise and diversification of major archaeal groups. BMC Biol 20, 154, doi:10.1186/s12915-022-01348-6 (2022).

53 Hug, L. A. et al. Community genomic analyses constrain the distribution of metabolic traits across the Chloroflexi phylum and indicate roles in sediment carbon cycling. Microbiome 1, 22, doi:10.1186/2049-2618-1-22 (2013).

54 Freches, A. & Fradinho, J. C. The biotechnological potential of the Chloroflexota phylum. Appl Environ Microbiol 90, e0175623, doi:10.1128/aem.01756-23 (2024).

55 Vinogradov, S. N. et al. A phylogenomic profile of globins. BMC Evol Biol 6, 31, doi:10.1186/1471-2148-6-31 (2006).

56 Gupta, K. J., Hebelstrup, K. H., Mur, L. A. & Igamberdiev, A. U. Plant hemoglobins: important players at the crossroads between oxygen and nitric oxide. FEBS Lett 585, 3843–3849, doi:10.1016/j.febslet.2011.10.036 (2011).

57 Vinogradov, S. N., Hoogewijs, D. & Arredondo-Peter, R. What are the origins and phylogeny of plant hemoglobins? Commun Integr Biol 4, 443–445, doi:10.4161/cib.15429 (2011).

58 De Visser, J. A. G. et al. Perspective: evolution and detection of genetic robustness. Evolution 57, 1959–1972 (2003).

59 Imachi, H. et al. Eukaryotes’ closest relatives are internally simple syntrophic archaea. bioRxiv, 2025.2002. 2026.640444 (2025).

60 Imachi, H. et al. Isolation of an archaeon at the prokaryote-eukaryote interface. Nature 577, 519–525, doi:10.1038/s41586-019-1916-6 (2020).

61 Yeates, T. O., Crowley, C. S. & Tanaka, S. Bacterial microcompartment organelles: protein shell structure and evolution. Annu Rev Biophys 39, 185–205, doi:10.1146/annurev.biophys.093008.131418 (2010).

62 Sutter, M., Melnicki, M. R., Schulz, F., Woyke, T. & Kerfeld, C. A. A catalog of the diversity and ubiquity of bacterial microcompartments. Nat Commun 12, 3809, doi:10.1038/s41467-021-24126-4 (2021).

63 Lyons, T. W., Reinhard, C. T. & Planavsky, N. J. The rise of oxygen in Earth’s early ocean and atmosphere. Nature 506, 307–315 (2014).

64 Edgar, R. C. MUSCLE: a multiple sequence alignment method with reduced time and space complexity. BMC bioinformatics 5, 1–19 (2004).

65 Capella-Gutiérrez, S., Silla-Martínez, J. M. & Gabaldón, T. trimAl: a tool for automated alignment trimming in large-scale phylogenetic analyses. Bioinformatics 25, 1972–1973 (2009).

66 Richter, O., Tao, J., Turba, A. & Ludwig, B. A cytochrome ba3 functions as a quinol oxidase in Paracoccus denitrificans. Purification, cloning, and sequence comparison. Journal of Biological Chemistry 269, 23079–23086 (1994).

67 Rieske, J. in Methods in enzymology Vol. 10 239-245 (Elsevier, 1967).

68 Lama, A., Pawaria, S. & Dikshit, K. L. Oxygen binding and NO scavenging properties of truncated hemoglobin, HbN, of Mycobacterium smegmatis. FEBS letters 580, 4031–4041 (2006).

69 Cline, S. W., Lam, W. L., Charlebois, R. L., Schalkwyk, L. C. & Doolittle, W. F. Transformation methods for halophilic archaebacteria. Canadian journal of microbiology 35, 148–152 (1989).

